# Dynamic model of bovine brucellosis to investigate control strategies in endemic settings

**DOI:** 10.1101/2022.03.14.483550

**Authors:** H. R. Holt, M. Walker, W. Beauvais, P. Kaur, J.S. Bedi, P. Mangtani, N.S. Sharma, J.PS. Gill, J. Godfroid, J. McGiven, J. Guitian

**Author notes:** Author contributions: HRH, MW, J. Godfroid, J. Guitian, PK, JSB, PM, NS, JPSG, JM designed the research; HRH, MW, WB, PK, JSB, J. Guitian performed the research; HRH and MW analysed data; HRH and J. Guitian wrote the paper; MW, J Godfroid and JM revised the paper. Author declaration The authors declare no conflict of interest. Funding: The project was enabled by joint funding from Biotechnology and Biological Sciences Research Council, UK (BB/L004836/1 & BB/L004895/1) and the Department of Biotechnology in India (BT/IN/Indo-UK/FADH/51/NSS/2013).

## Abstract

Brucellosis imposes substantial impacts on livestock production and public health worldwide. A stochastic, age-structured model incorporating herd demographics was developed describing within- and between-herd transmission of *Brucella abortus* in dairy cattle herds. The model was fitted to data from a cross-sectional study conducted in Punjab State of India and used to evaluate the effectiveness of control strategies under consideration. Based on model results, stakeholder acceptance and constraints regarding vaccine supply, vaccination of replacement calves in large farms should be prioritised. Test and removal applied at early stages of the control programme where seroprevalence is high would not constitute an effective use of resources. Critically, under current model assumptions, significant numbers of animals ‘removed’ (culled or not used for breeding) in this strategy would be removed based on false positive results. To achieve sustained reductions in brucellosis, policymakers must commit to maintaining vaccination in the long term, which may eventually reduce frequency of infection in the livestock reservoir to a low enough level for elimination to be a realistic objective. This exercise provided important insights into the control of brucellosis in India, which has the largest cattle population globally, and a general framework for evaluating control strategies in endemic settings.

## Introduction

Brucellosis is a bacterial zoonosis imposing significant impacts on human health, livestock production and international trade of livestock products (1,2). In 2011, the World Bank ranked brucellosis among the top ten diseases in cattle, buffalo, sheep and goats and camelidae in terms of ‘Livestock Units Lost’ (2). The main transmission routes for human infection are foodborne, as a result of consumption of raw milk or unpasteurized dairy products, and direct contact with contaminated tissues (placenta, aborted foetuses, carcasses) and parturition fluids from infected livestock. Brucellosis causes acute febrile illness which, if not diagnosed and treated, can lead to chronic debilitation (2–4). Targeting the livestock reservoir through vaccination, culling of animals deemed to be infected and movement restrictions have led to elimination of the disease in several countries (5,6). However, these strategies require significant resources and long-term commitment and are difficult to implement in areas where smallholder systems predominate. As a result, the disease remains endemic in many countries of Africa, Asia, Latin America and the Middle East where it is one of the major contributors to disability-adjusted life years (DALYs) lost as a result of foodborne illness (3,7–9).

India is the world’s leading milk producer, with the world’s largest bovine population that are kept predominantly in smallholder systems. Bovine brucellosis is considered endemic throughout the subcontinent (7,10–13), however, high quality epidemiological studies are lacking (14). India is also home to approximately 18% of the world’s human population and it is therefore likely that a significant proportion of the global burden of brucellosis infections in both people and cattle occurs here (15,16). Control of brucellosis, and other cattle diseases, is particularly complex in India, where slaughter of cattle is not usually acceptable and is illegal in several states (11,14). Brucellosis control has gained interest from Indian policymakers in recent years; a pilot programme termed the ‘*Brucella* free village’ was launched in 2016; aiming to eliminate brucellosis within 50 selected villages in 10 states (17). This programme originally considered testing all adult livestock in selected villages, vaccinating those testing seronegative and segregating those testing seropositive to create disease free zones (17). In addition, a ‘National Animal Disease Control Programme for Food and Mouth Disease (FMD) and Brucellosis’ was approved by the Cabinet of the Central Government on the 31st May 2019 and is currently in the planning stages. The National Control Programme proposes vaccination as the primary control strategy, however, one of the major challenges in the National Control Programme is the procurement of sufficient vaccine doses. It is unlikely that very high coverage of the vaccine will be achieved in the short-term, given current low availability, logistics of implementing large-scale vaccination campaigns, reluctance of some farmers to vaccinate as live vaccines may induce abortion in livestock and the sheer size of the bovine population to be vaccinated (7,18). Therefore, evidence upon which the vaccination effort could be targeted and whether vaccination at low coverage can still be effective are particularly timely.

In order to evaluate the likely effectiveness of competing intervention strategies, we propose a stochastic model for bovine brucellosis due to *Brucella abortus*, which captures both herd demographics and disease dynamics over time. The model is parameterized using data from a recent cross-sectional study conducted in 425 dairy farms in Punjab state, which is the state that produces the most cattle and buffalo milk per capita in India (12). Previous work has demonstrated that brucellosis is widespread in dairy farms in Punjab where exposure of people in direct contact with cattle and buffaloes is common (4,19,20). Control strategies simulated were those considered by Indian brucellosis control programmes. The effect of vaccinating varying proportion of herds, and animals within herds, with (*Brucella* free village) and without (National Control Programme) implementation of test and removal at the inception of the control programme was investigated. To our knowledge, this is the first brucellosis model that explicitly incorporates within- and between-herd transmission in a setting in which the control of programme takes place without movement restrictions, which are difficult to implement in most endemic settings.

## Materials and Methods

### Disease transmission model

A dynamic age-structured stochastic model incorporating dairy herd demographics was written in R (21) to capture transmission of *B. abortus* within and between cattle herds. Transition between compartments follows a random Poisson process using the event-driven Gillespie stochastic simulation algorithm (SSA) implemented using the R package GillespieSSA (22). The model tracks numbers of individual cattle in each age group and state; susceptible, infected (exposed), infectious and vaccinated (Figure 1. & Supplementary Information).

**Figure 1:**
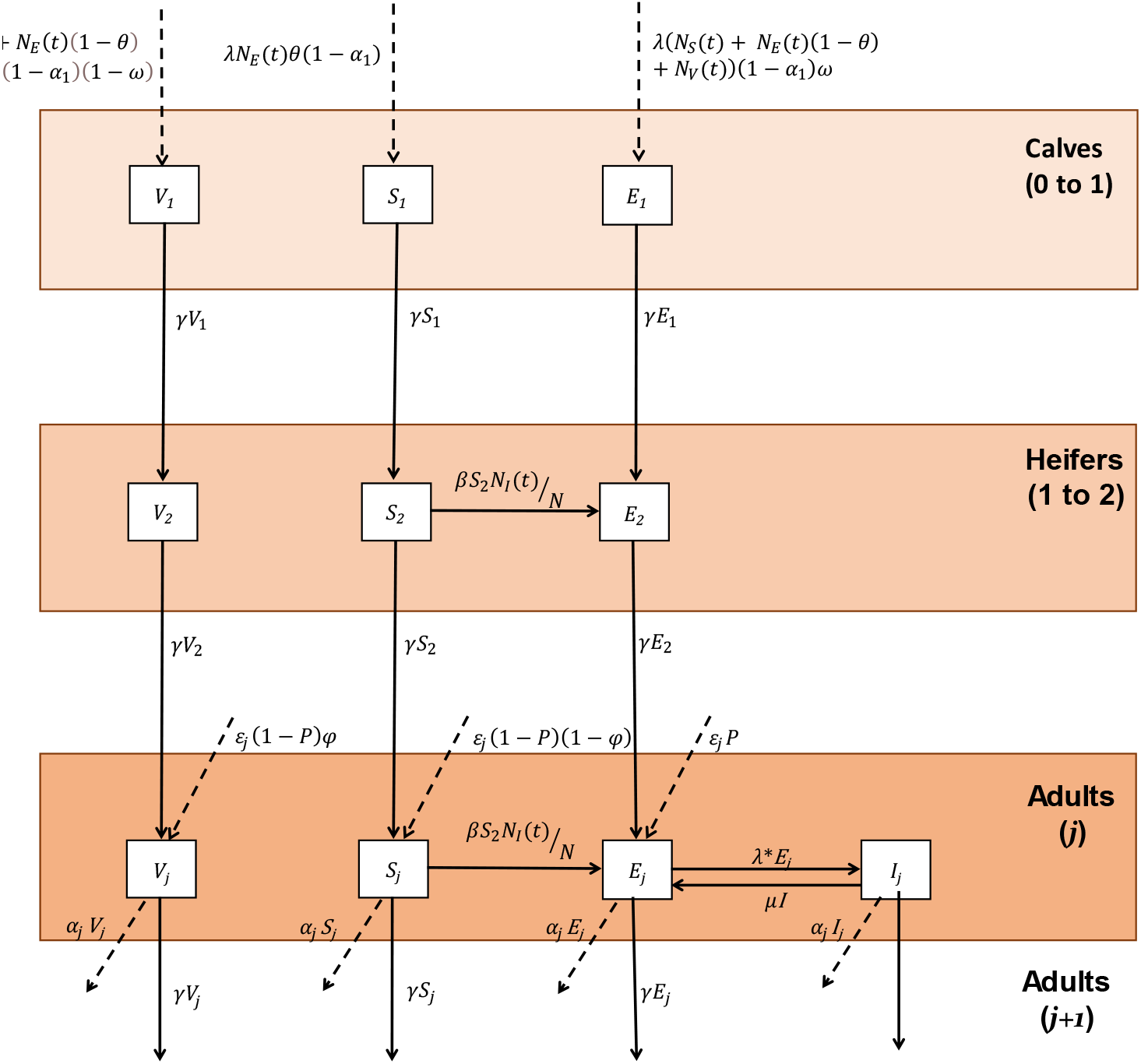
Conceptual model framework showing the transition between different compartments, where 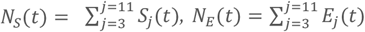 and 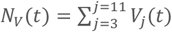 denote, respectively, the total number of susceptible, exposed and vaccinated adults in the herd.

### Model Events

Replacement animals are modelled as being sourced either from new births on the farm, with λ denoting the annual birth rate, or from the purchase of new animals with rate parameter *ε*_*j*_. Male calves, new-borns sold off the farm or calves that die in their first year of life cannot become infectious whilst on the farm and are therefore not included in the model; these events are incorporated in a single rate parameter, *α*_1_. *I*n other age groups, livestock die, are sold or are no longer reproductively active described by a single rate parameter, *a*_*j*_.

Within herds, both horizontal and vertical transmission are captured. Calves born to infected cows are either susceptible, *S*_*1*_, or infected (exposed), *E*_*1*_, vertical transmission from mother to calf occurs at rate *θ*. Infected cattle infect susceptible cattle when they abort or give birth. Following a period of infectiousness, where *B. abortus* organisms can survive in the environment, cattle return to the infected (exposed) compartment at rate *µ* until they give birth again. Transmission of *Brucella* spp. is via proximity to calving livestock, contaminated environments, *in utero* or during service/artificial insemination. Therefore, as herds in Punjab are primarily sedentary and effective contacts are unlikely to occur between livestock from different farms (12), it was assumed that between-herd transmission only occurs through purchase of infected cattle, which is a function of the number of annual purchases (*ε*_*j*_) and the likelihood a replacement animal is infected (current animal-level prevalence; *P*).

### Model parameterisation

The model was parameterised using data from a sero-survey of 425 dairy herds in rural Ludhiana district of Punjab between 2015 and 2017; 139 herds had at least one positive animal (median within-herd prevalence 33.3%) (18). This was supplemented with information collected from a survey of 409 household herds (20) and published literature (Supplementary Information). Two zero-inflated negative binomial distributions for purchase rates were fitted using maximum likelihood estimation in the package fitdistrplus (23), one for herds with less than nine adult females and one for herds with nine or more based on observed data that distributions of purchases. The effective contact rate within a farm (‘beta transmission’ parameter) was fitted using ABC techniques. See Supplementary Information and Holt et. al., (2021) for further details.

### Simulation of control scenarios

Simulations of the impact of different control scenarios based on vaccination are performed at village level, as this is the unit considered for control by the *Brucella* free village campaign and National Control Program. Initial herd sizes for 180 dairy herds per village were sampled from zero-truncated fitted distribution and the model was run for 75 years to stabilise. The model was used to investigate the following:

1. <u>Vaccination strategies:</u>
  i. the effect of vaccinating different subsets of herds (**A:** all herds, **B:** 50% of herds (randomly selected), **C:** large herds only (9+ animals)
  ii. the effect of vaccinating different proportions of all livestock in the herd at the inception of the control programme (year 1)
  iii. the effect of vaccinating different proportions of calves annually
2. <u>‘Test and removal’:</u> Selected vaccination strategies were simulated with and without serological testing of all animals and isolation of those found to be positive at the inception of the control programme (as proposed in the *Brucella* free village programme).

The disease transmission model (Figure 1) tracks the numbers of livestock in each state; susceptible (*S*_*j*_), infected (*E*_*j*_), vaccinated (*V*_*j*_) (all age groups) and infectious (*I*_*j*_; adults only), as well as vaccine doses and the number of animals ‘removed’ from the dairy herds in test and removal strategies. Targets for control were set using prevalence (percentage of infected animals in a village), as this is measurable via surveillance. Here we define ‘control’ as animal-level prevalence below 1%, it was assumed that at this point Punjab could consider moving towards elimination. Predicted (cumulative) incidence, the percentage of animals infected with *B. abortus* per year, is also presented.

## Results

### Scenarios without ‘test and removal’ policy

Despite there being no new infections (incidence = 0%) in some of the scenarios after running the control programme for 30 years (Figure 2A), median animal prevalence of *B. abortus* only reached 0% in the highest coverage Scenario A strategies, which involved vaccinating 100% of replacement calves (Figure 2B). This is due to the long lifespan of Indian cattle and the assumption that infected cows can remain infected and infectious for their reproductive life. Control strategies which included annual vaccination within all herds in a village (Scenario A), reduced the median animal level prevalence from 12.9% (year 0) to below 1% within 30 years (Figure 2A) in scenarios where 75% or 100% of calves were vaccinated.

**Figure 2.**
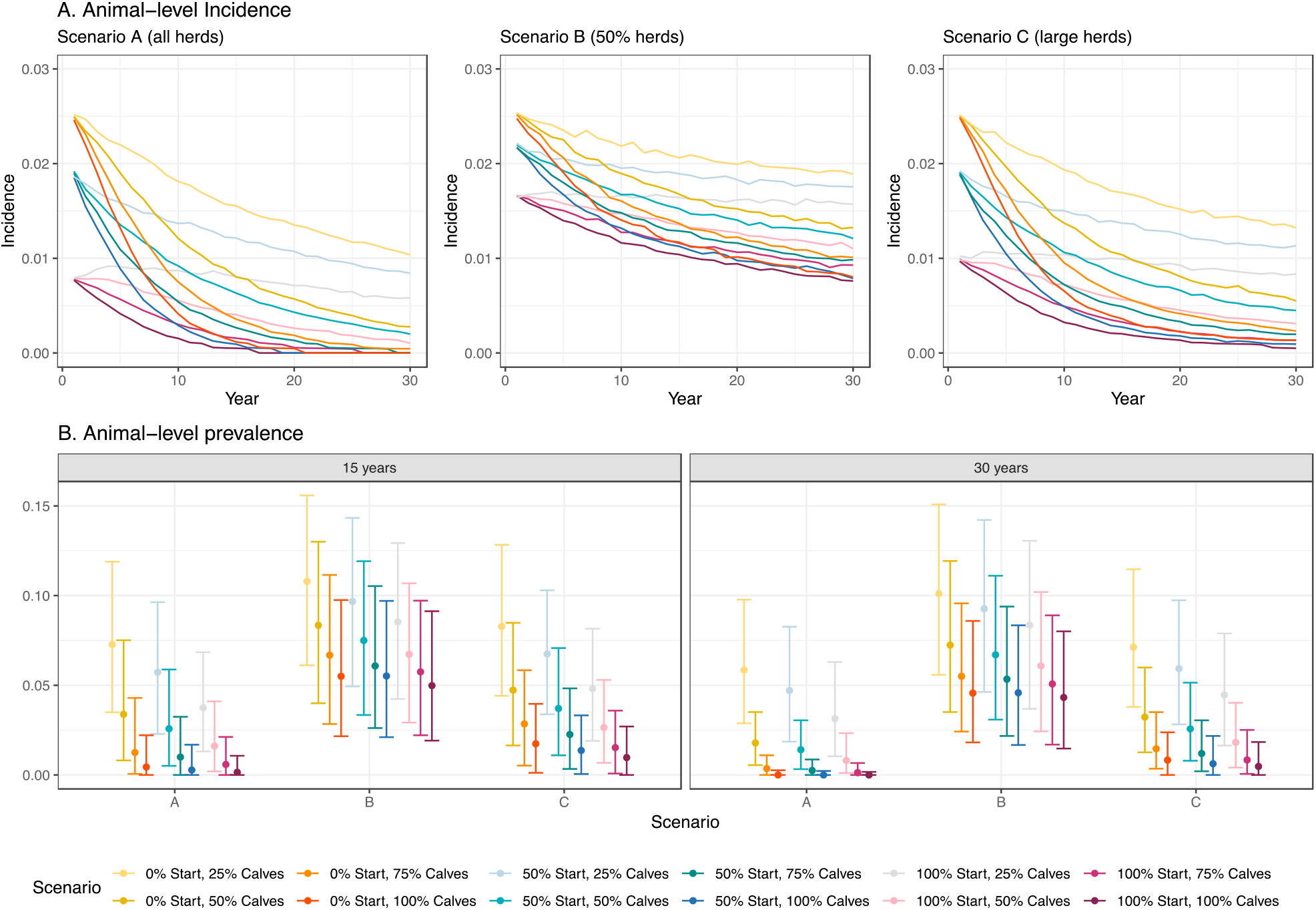
Incidence (proportion of animals infected per year) from year 1 of the control programme (A); proportion of infected bovines after employing each strategy for 15 and 30 years (B), according to each strategy employed - based on 1000 iterations/villages (‘Start’ = % of the herd (all ages) vaccinated in year 1 of the control programme and ‘Calves’ refers to the % of calves vaccinated)

Vaccination of animals in herds with nine or more animals (Scenario C), where 100% or 75% of calves were vaccinated annually resulted in control with at least 75% probability (i.e. percentage of iterations, Figure 3A). Control was not achieved in any of the strategies where 50% of herds were chosen at random to be enrolled in the vaccination programme (Scenario B) and varying proportions of calves within these herds are vaccinated annually (>80% of iterations). The model projected that Scenarios A and C would avert the most bovine infections, with a median of over 1000 infections averted per village during the 30-year control period when 75% or 100% of replacement calves were vaccinated (Figure 3B). Employing Scenario C, as opposed to Scenario A, was estimated to delay control by a median of 5 years (Figure 3C). However, Scenario C reduces the vaccine doses by around a third and saves additional resources as only 35% of farms need to be vaccinated as opposed to 100% (Figure 3D). Scenario C also resulted in a bigger reduction in the prevalence of the disease compared to annual vaccination of 50% of herds (Scenario B).

**Figure 3:**
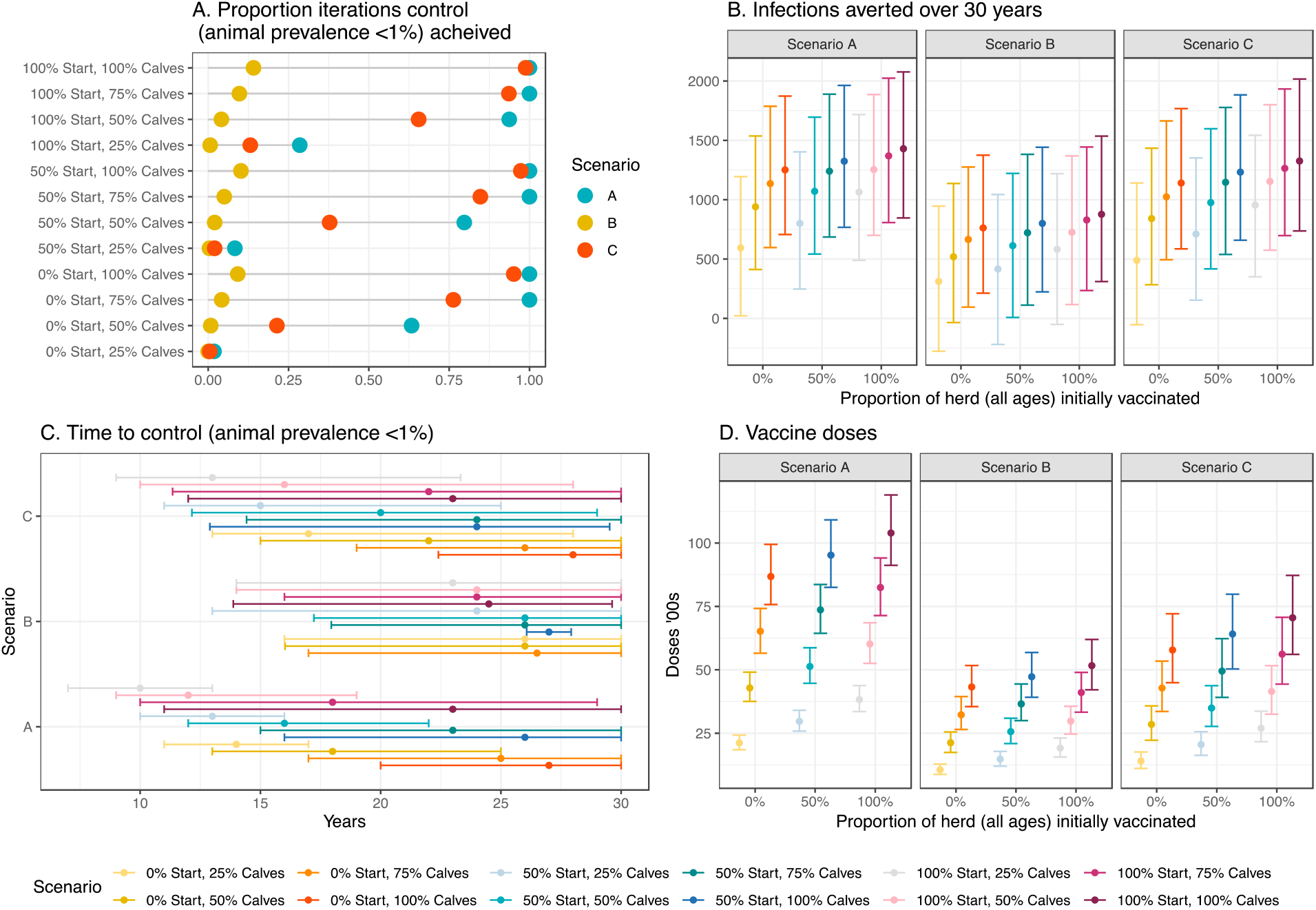
Proportion of times ‘control’ (defined as animal level prevalence below 1%) is achieved (A), number of infections averted per village (compared to baseline scenario of no control) (B), time to achieve control (C) and number of vaccine doses (D) and after employing each strategy for 30 years, based on 1000 iterations/villages (‘Start’ = % of the herd (all ages) vaccinated in year 1 of the control programme and ‘Calves’ refers to the % of calves vaccinated. Median and 95% Credible Interval are presented in A, B, & D.

### Scenarios with ‘test and removal’ policy

The impact of combining vaccination of calves within all herds (Scenario A) and large herds (Scenario C) with testing of adults at the inception of the control programme and removing those giving a positive serological result from the farm was also investigated. These simulations resulted in control (animal-level prevalence <1%) in more than 85% of villages/iterations (Figure 4D), within a median of 1 year (Figure 4E). However, under the conservative assumption of a test sensitivity and specificity of 90%, these scenarios did not remove all infected livestock in year 1 (Figure 4A) and would result in the removal of large numbers of uninfected animals (false positives). In these scenarios a median of 102 (95% Credible Interval (CI): 82; 132) uninfected and 141 (95% CI; 91; 207) infected animals were removed from population per village (around 20% village population; 7 uninfected animals per 10 infected removed). Over the 30-year period, the number of new infections averted per village ranged between 1197 in Scenario C, where 25% of replacement calves were vaccinated, to 1393 in Scenario A, where 100% of calves were vaccinated. Compared to the equivalent vaccination-only scenarios, the addition of test and removal resulted in substantial reductions in new cases in scenarios with low vaccination coverage (807 fewer infections in Scenario C with 25% of replacements vaccinated). However, in scenarios with higher coverage similar reductions in new infections were achieved (142 fewer infections in Scenario A with 100% of replacements vaccinated) (Figure 4).

**Figure 4:**
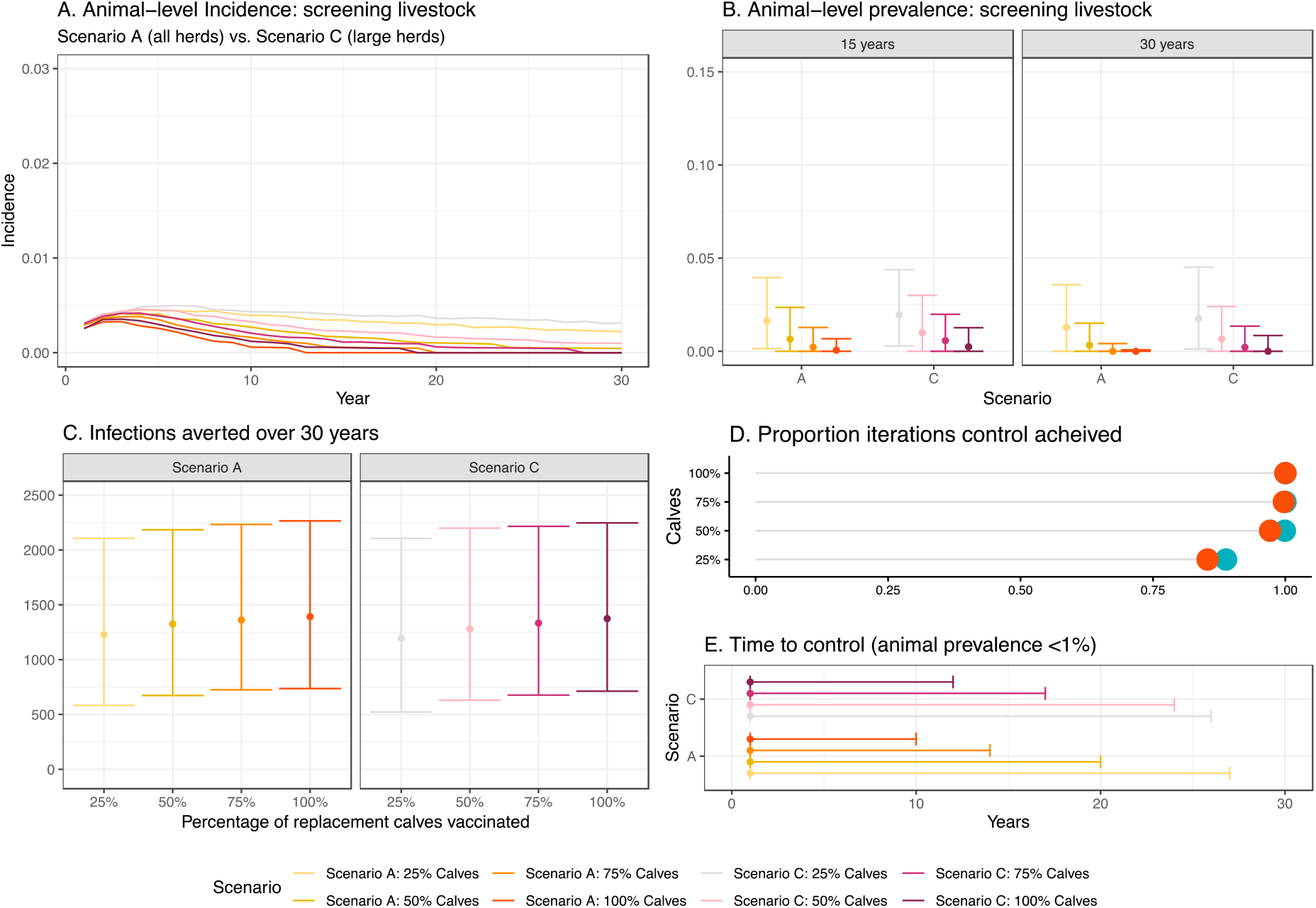
Cumulative incidence over time according to strategy employed when seropositive animals are removed in year 1 (A), Prevalence (proportion of infected bovines) after employing each strategy for 15 and 30 years (B), number of infections averted per village (compared to baseline scenario of no control) (C), proportion of times ‘‘control’ (defined as animal level prevalence below 1%) is achieved (D) and time to achieve control in a village (E). In all scenarios test and removal is performed in year 1 and model is run over 30 years for 1000 iterations/villages ‘Calves’ refers to the % of calves vaccinated. Median and 95% Credible Interval are presented in B, C & E.

## Discussion

Despite successful elimination of bovine brucellosis in some countries, its control in many settings, including India, remains elusive (7). Lack of management options for positive animals is often perceived as one of the main barriers to controlling brucellosis in India (18,24). However, this study demonstrated that control (defined here as animal level prevalence below 1%) of *Brucella* infections in bovines in endemic areas could be achieved through vaccination, without the need to vaccinate all animals or implement a ‘test and removal’ policy. Although control, as opposed to elimination, will not result in the attainment of Official Brucellosis Free (OBF) status, a marked decrease in the incidence of infection can be expected to result in comparable reductions in financial losses to farmers and incidence of human infection (25–32).

Sustainability, due to limited resources and political will, has constrained brucellosis vaccination campaigns in many contexts (5,33). Due to the chronic nature of the disease and the long lifespan of cattle in India, if vaccination campaigns are to be an effective use of resources the commitment must be long term, the goal (e.g. sustained reduction/control vs. elimination) should be achievable and the control strategy realistic. The results suggest calfhood vaccination is likely be successful at lowering the animal prevalence of the infection below 1% under the current assumptions regarding model parameters. However, even with the most aggressive vaccination strategy (vaccination of 100% of animals at the start of the control program and 100% of calves annually) the median time estimated to achieve this was 10 years, with 95% of estimates falling between 8 and 14 years. The National Control Program, which envisages 100% vaccination of female cattle and buffalo calves once in their lifetime, has initially been budgeted for 5 years (17). However, the model projected that it would take a median of 14 years to achieve control when only calves are vaccinated. Furthermore, vaccination is unlikely to result in elimination, therefore calfhood vaccination would need to be implemented indefinitely to prevent re-emergence of brucellosis (5).

Strategies applied to complex settings require adaptation and should consider the diversity of the livestock systems, infrastructure and available resources. A lesson to be learnt from previous brucellosis control programs in other countries is that test and slaughter should not be introduced too early, without the resources for compensation to farmers, or where the baseline prevalence is so high that a large proportion of the livestock population requires culling (3,5,34). For example, in the European Union, legislation related to the control and eradication programs of bovine brucellosis started in 1962, and only seven out of fifteen Member States were recognized as OBF in 1999, 37 years after inception (35). In India, the *Brucella* free village campaign initially included a component on test and removal of infected livestock and if this was implemented then elimination would be accelerated with animal prevalence projected to go below 1% within the first year. However, given that 12.9% (95% CI: 9.2; 17.6) of animals in the study area were estimated to be seropositive at the start of the control programme, huge resources would be required to identify these animals and remove them from the population, especially since the *Brucella* free village programme proposes to maintain infected cows in an area separate from the village (17). In addition, replacement livestock would need to be sourced externally which may result in infected livestock being re-introduced in the village (36). Alternatively, if sustained reductions in prevalence were achieved through targeted vaccination then re-allocation of resources towards elimination activities in certain farms, villages, states – or at a national level may become feasible.

Stakeholder consultation revealed that vaccine availability is a limiting factor and one of the major activities of the National Control Program will be vaccine procurement (18). The situation is likely to become even more challenging as Indian Immunologicals who introduced brucellosis vaccine – ‘Bruvax’ to the Indian market and are the primary manufacturer of the vaccine are now engaging with SARS-CoV-2 vaccine production (18,37,38). The SARS-CoV-2 pandemic has also disrupted supply chains for reagents and consumables, resulting in unpredictable shutdowns of manufacturing and movement restrictions (39,40). Therefore, if pursued, policymakers should ensure brucellosis vaccine programs are resilient to such shocks prior to implementation. In addition to the difficulties and costs associated with the manufacture and procurement of the vaccine, there are logistical issues surrounding its deployment.

The most effective strategy for brucellosis control may not comprise the most efficient use of resources (41); policymakers can use the information presented here to assess which vaccination scenarios will be optimal based on available doses and resources. For example, vaccinating 100% of calves under Scenario C (targeting large herds only) and Scenario A (all herds) with no test and removal and no initial vaccination was estimated to save similar numbers of infections per village (median 1326 vs 1430 over 30 years) and resulted in the control of *Brucella* spp. (animal-level prevalence < 1%) in 100% of iterations/villages. However, the former required around 30% fewer vaccine doses and only animals within around a third of herds to be vaccinated. As well as increased herd sizes, this finding is driven by the higher rate of purchasing new livestock in these farms, therefore screening of animals prior to sale may also present a potential future control option; the model has already demonstrated that this can be a useful control strategy at farm-level (18). Stakeholders envisioned calfhood vaccination being more acceptable to farmers and veterinary offices due the negative effects of the vaccine that can occur in adult livestock and the potential to reduce needlestick or other injuries (18). They also reported that larger commercialized farms would be easier to target due to their heightened awareness of the disease and willingness to invest in herd health.

The aim of this study was to predict the *relative impact* of different control scenarios; hence the parameters were mostly fixed in order to maximise comparability. However, there is uncertainty surrounding some of the parameter estimates, particularly the probability a calf born to an infected cow is infected (*PIC*). This was explored in sensitivity analysis using a worst case (20%) estimate and this did not alter the relative effectiveness of the different scenarios (18). A conservative estimate for test performance (90% sensitivity and specificity, Rose Bengal Test) was used, however, it is likely that accuracy, particularly specificity, is higher which may have led to an overestimate of the number of susceptible animals giving false positive result (18). A vaccine efficacy of 80% was selected, however, a recent systematic review and meta-analysis indicates this may be optimistic (42). In this study, S19 at a dose of 10^9^ colony forming units (CFUs) was estimated to be 75% (95% CI: 48 – 88%) and 72% (95% CI: 30.9 to 84%) efficacious against abortion and infection, respectively (42). The study by de Oliveira et al., (2021) found that a dose of 10^9^ CFU had the highest vaccine efficacy which is 50-80 times lower than the dose currently recommended by the OIE for subcutaneous administration. If this dose was shown to be efficacious in this setting this could also help address the issue of lack of vaccine doses.

The model already includes several components omitted in previous simulation models of control strategies, allowing for stochastic extinction within a herd and contact rates between livestock to vary by explicitly modelling within- and between-herd transmission. A novel method of the seeding of infected livestock through trade is used and infected animals become infectious following a calving/abortion event, as opposed to at a fixed rate. This results in more realistic model simulations making the outputs more applicable to the issue of sustainable brucellosis control in complex settings. Potential further applications of the model include simulation of additional control strategies, including separation of cattle and buffalo in the model to simulate tailored interventions, testing model assumptions including test performance and vaccine efficacy, and adaptation to other settings or diseases of livestock for which control relies on combining vaccination and test-and-slaughter and for which within- and between-herd transmission should be explicitly simulated.

In conclusion, taking Punjab State of India as a case study, this work demonstrates the application of a stochastic mathematical model to gain important insights into the control of bovine brucellosis, among the livestock zoonoses responsible for a high global burden (2,43). The results suggest that, under current model assumptions and parameters, control can be achieved through targeted vaccination without the need to achieve perfect coverage. Targeting animals within large herds and replacement calves to receive vaccination may be a more efficient use of resources than blanket vaccination, which is not feasible while vaccine doses are limited. However, in order for reductions to be sustained, policymakers must commit to long-term vaccination as control is unlikely to be achieved within the proposed timeframe of the current programs. In the absence of additional control measures, it would be necessary to pursue vaccination indefinitely. It is imperative that stakeholders are aware of this before vaccination campaigns are implemented. Test and removal of livestock during year 1, although being efficient in reducing animal prevalence, is unlikely to be an acceptable strategy for livestock owners or an effective use of resources at this stage in the control program in Punjab.

## Acknowledgements

The authors are grateful to dairy sector stakeholders for sharing their views with the project team. They also express thanks to Dr. Amanda Minter for providing advice on model fitting and to Dr. Tom Sumner for their constructive comments.

