## Supplementary Information for "Dynamic model of bovine brucellosis to investigate control strategies in endemic settings"

**This PDF file includes:**

Supplementary text
Figure S1
Table S1
SI References

**Supplementary Information Text**

**Model definition**

The model is structured into 11 age groups, denoted by subscript $j$, comprising ten 1-year intervals from 0 to 10 (0-1, 1-2, 2-3, etc.) and an additional group for animals aged >10 years. Only females are included in the model. Within each age group, animals can be either susceptible $S$, exposed/infected, $E$ or vaccinated, $V$. Additionally, animals aged > 2 years (i.e. $j = 3 \ldots 11$)—here defined as adults—already exposed can become infectious, $I$, for a period $1/\mu$ after giving birth. In the following description all parameters are defined in Table S1, as well as in the text. We begin by defining the probabilities associated with changes in the number of susceptible, exposed and vaccinated calves, $j = 1$, at time $t$,

$$P[S_1(t+\delta t) = S_1(t) + 1] = \delta t[\lambda(N_S(t) + \; N_E(t)(1-\theta) + N_V(t))(1-\alpha_1)(1-\omega)]$$

$$P[S_1(t+\delta t) = S_1(t) - 1] = \delta t\ [\gamma S_1]$$

$$P[E_1(t+\delta t) = E_1(t) + 1] = \ \delta t[\lambda N_E(t)\theta(1-\alpha_1)]$$

$$P[E_1(t+\delta t) = E_1(t) - 1] = \ \delta t\ [\gamma E_1]$$

$$P[V_1(t+\delta t) = V_1(t) + 1] = \ \delta t[\lambda(N_S(t) + \; N_E(t)(1-\theta) + N_V(t))(1-\delta_1)\omega]$$

$$P[V_1(t+\delta t) = V_1(t) - 1] = \ \delta t\ [\gamma V_1] \quad (1)$$

Here, $N_S(t) = \sum_{j=3}^{j=11} S_j(t)$, $N_E(t) = \sum_{j=3}^{j=11} E_j(t)$ and $N_V(t) = \sum_{j=3}^{j=11} V_j(t)$ denote, respectively, the total number of susceptible, exposed and vaccinated adults in the herd. Parameter $\lambda$ denotes the per capita birth rate (also the rate at which exposed adults become infectious), $\theta$ is the probability of maternal transmission from an exposed adult, $\alpha_1$ is the rate of removal of newborn calves (because of death or immediate sale) and $\omega$ is the probability that a calf is vaccinated. Parameter $\gamma$ denotes the transition rate from one age group to the next.

Probabilities defining transitions in the number of heifers, $j = 2$, in each state at time $t$ are as follows:

$$P[S_2(t+\delta t) = S_2(t) + 1] = \delta t\ [\gamma S_1]$$

$$P[S_2(t+\delta t) = S_2(t) - 1] = \delta t\ \left[\left(\gamma S_2 + \frac{\beta S_2 N_I(t)}{N}\right)\right]$$

$$P[E_2(t+\delta t) = E_2(t) + 1] = \ \delta t\left[\left(\gamma E_1 + \frac{\beta S_2 N_I(t)}{N}\right)\right]$$

$$P[E_2(t+\delta t) = E_2(t) - 1] = \ \delta t\ [\gamma E_2]$$

$$P[V_2(t+\delta t) = V_2(t) + 1] = \ \delta t\ [\gamma V_1]$$

$$P[V_2(t+\delta t) = V_2(t) - 1] = \delta t\,[\gamma V_2] \tag{2}$$

where $N_I(t) = \sum_{j=3}^{j=11} I_j(t)$ is the total number of infectious adults, $N$ is the herd size, and $\beta$ is the per capita contact rate among animals. Note here that the contact rate is assumed to be *independent* of herd size.

Finally, event probabilities for adults, $j = 3 \dots 11$, are given by:

$$P\left[S_j(t+\delta t) = S_j(t) + 1\right] = \delta t\,\left[\gamma S_{j-1} + \varepsilon_j(1-P)(1-\varphi)\right]$$

$$P\left[S_j(t+\delta t) = S_j(t) - 1\right] = \delta t\,\left[\left(S_j(\alpha_j + \gamma) + \frac{\beta S_j N_I(t)}{N}\right)\right]$$

$$P\left[E_j(t+\delta t) = E_j(t) + 1\right] = \delta t\left[\gamma E_{j-1} + \frac{\beta S_j N_I(t)}{N} + \varepsilon_j P\right]$$

$$P\left[E_j(t+\delta t) = E_j(t) - 1\right] = \delta t\,\left[E_j\left(\alpha_j + \gamma + \lambda\right)\right]$$

$$P\left[I_j(t+\delta t) = I_j(t) + 1\right] = \delta t\,\left[\gamma I_{j-1} + \lambda E_j\right]$$

$$P\left[I_j(t+\delta t) = I_j(t) - 1\right] = \delta t\left[I_j\left(\alpha_j + \gamma + \mu\right)\right]$$

$$P\left[V_j(t+\delta t) = V_j(t) + 1\right] = \delta t\,\left[\gamma V_j + \varepsilon_j(1-P)\varphi\right]$$

$$P\left[V_j(t+\delta t) = V_j(t) - 1\right] = \delta t\,\left[V_j\left(\alpha_j + \gamma\right)\right] \tag{3}$$

where $\varepsilon_j$ is the rate of incoming newly purchased animals which takes a positive value sampled from one of two negative binomial distributions, dependent on herd size (fitted to observed purchase data) for $j = 4$ and 0 otherwise, under the assumption that all purchased animals are in the 3-4 age group, $P$ is the prevalence of infection (i.e., exposed animals) among *all* simulated herds. The terms $\varepsilon_j(1-P)(1-\varphi)$, $\varepsilon_j P$ and $\varepsilon_j(1-P)\varphi$ thus determine the number of incoming animals that are either susceptible, exposed or vaccinated, as a function of the prevalence of infection across all herds. Parameter $\varphi$ donates the probability a new purchase is vaccinated, this is updated after every 12 months of simulation to reflect the changing proportion of animals in the village that are vaccinated. Parameter $\delta_j$ defines the rate of sale or death (combined) of adults which takes a value 12/72 for $j = 3 \dots 10$ and 0.84 for $j = 11$. When implementing vaccination, immunity is assumed to be lifelong.

In order to investigate the impact of different control scenarios, farms within villages were simulated. The average population of a village in the study area is approximately 1500 people with an average household size of 5.2 persons (1) and 61.0% of families are keeping cattle (2). Therefore, it was assumed an average village would contain 180 herds. Median herd size in the study area was four (2.5$^{th}$; 97.5$^{th}$ percentile; 1; 16) with a maximum of 24 animals (3). A negative binomial distribution was fitted to existing herd size data and initial herd size sampled from zero-truncated fitted distribution. The 180 farms were simulated for 1000 iterations (1000 villages) for 75 years in yearly increments using the same values as the within-herd model in order to reach endemic stability. Initial herd size was sampled from the zero-truncated negative binomial distribution and initially all cattle were in the age four compartment to ensure age distribution within herds was similar to observed data after running the model to endemic stability.

**Table S1.** Model parameters definition and values

| Parameter | Definitions | Value | Source |
|---|---|---|---|
| $\lambda$ | Calving rate (and rate that exposed animals become infectious) | 0.605 per year | (3) |
| $\gamma$ | Transition rate between age groups | 1 per year | |
| $\varepsilon_j$ | Rate of purchasing new animals | Sampled for $j$ = 4; 0 otherwise | Fitted to observed data from (3) |
| $\alpha_1$ | Rate of removal of newborn calves (composite parameter capturing death and sale of calves) | 0.685 per year | (3) |
| $\alpha_j$ | Rate of removal of adults (composite parameter capturing death, sale and end of reproductive activity of adults) | 12/72 for $j = 3 \ldots 10$<br>0.835 for $j = 11$ | Estimated from age distribution in (3) |
| $\beta$ | Cattle-to-cattle effective contact rate | 5.279 | This work |
| $\mu$ | Rate of loss of infectiousness (composite parameter capturing period bacteria are shed plus their survival in the environment) | 12/4 per year | (4,5) |
| $\theta$ | Probability calf born to infected animal is infected | 0.05<br>(0.2 in sensitivity analysis) | (6–8) |
| $\omega$ | Proportion of calves that are vaccinated | Varied value | 0.25; 0.5; 0.75 or 1.0 depending on scenario |
| $\varphi$ | Probability that a purchased animal is vaccinated | Varied value | Initially 0, updated annually when simulating control strategies |
| $VE$ | Vaccine efficacy | 0.8 | (9) |
| $Se$ | Test sensitivity | 0.9 | (10) |
| $Sp$ | Test specificity | 0.9 | |
| $Buf$ | Proportion of animals that are buffalo (vs. cows) | 0.45 | (3) & Supplementary Information |

**Fitting Beta transmission parameter**

The rate that susceptible cattle become infected is driven by the effective contact rate ($\beta$), which was fitted to the observed within-herd seroprevalence data presented in Holt et al, (2021) using an ABC algorithm. The algorithm samples a value of $\beta$ from a prior distribution and then produces a stochastic simulation for each of the 425 observed herds using the model function. An uninformative uniform prior bounded between 1 and 10 was used to describe beta and 5000 beta values were sampled from this distribution. For each value of beta, each of the 425 herds was simulated for 75 years in yearly increments. Initial herd sizes for each simulated herd were set to match the corresponding observed herd and all animals within herds were initially negative until an infected animal was purchased. The goodness of fit metric used for the model was the sum of the difference between the observed and simulated mean and standard deviation of the within herd prevalence (Fig. S1). Values of beta where the difference between the observed and simulated data was less than 0.005 were retained. Rather than a posterior distribution, the median value of Beta (5.279) was selected as the objective of subsequent model simulations was to compare the relative impact of different control strategies, a fixed value would reduce the variability between the different scenarios to make the results more comparable. However, the posterior distribution could be used in future simulation exercises.

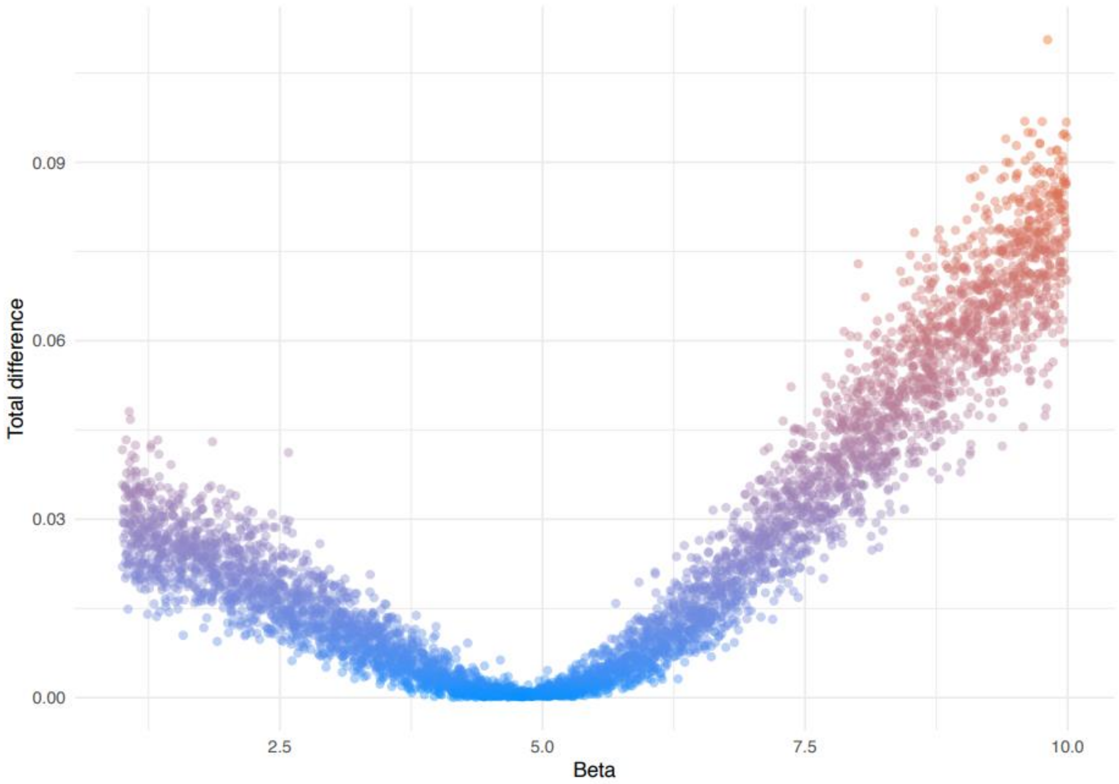

*Fig. S1 Scatterplot showing the total difference between the observed and simulated herd for each value of beta. The total difference was calculated as the sum of the difference between the observed and simulated mean and the observed and simulated standard deviation of the within-herd prevalence*
